# The proteotranscriptomic characterization of venom in the white seafan *Eunicella singularis* elucidates the evolution of Octocorallia arsenal

**DOI:** 10.1101/2024.05.31.596435

**Authors:** Maria Vittoria Modica, Serena Leone, Marco Gerdol, Samuele Greco, Didier Aurelle, Marco Oliverio, Giulia Fassio, Khadija El Koulali, Célia Barrachina, Sebastien Dutertre

## Abstract

All the members of the phylum Cnidaria are characterized by the production of venom in specialized structures, the nematocysts. Venom of jellyfish (Medusozoa) and sea anemones (Anthozoa) has been investigated since the 1970s, revealing a remarkable molecular diversity. Specifically, sea anemones harbour a rich repertoire of neurotoxic peptides, some of which have been developed in drug leads. However, venoms of the vast majority of Anthozoa species remain uncharacterized, particularly in the class Octocorallia. To fill this gap, we applied a proteo-transcriptomic approach to investigate the venom composition in *Eunicella singularis*, a gorgonian species common in Mediterranean hard-bottom benthic communities. Our results highlighted the peculiarities of the venom of *E. singularis* with respect to sea anemones, which is reflected in the presence of several toxins with novel folds, worthy of functional characterization. A comparative genomic survey across the octocoral radiation allowed us to generalize these findings and provided insights into the evolutionary history, molecular diversification patterns and putative adaptive roles of venom toxins. A comparison of whole-body and nematocyst proteomes revealed the presence of different cytolytic toxins inside and outside the nematocysts. Two instances of differential maturation patterns of toxin precursors were also identified, highlighting the intricate regulatory pathways underlying toxin expression.

## Introduction

Cnidaria is the most ancient venomous lineage among the extant Metazoa, having originated 600-800 million years ago [1,2]. Venom is produced in the nematocysts, specialized cells with high morphological diversity [3,4] and is involved in different functions including prey capture, defence from predators, and possibly also *intra-* and *inter-*specific competition [5]. In addition, in sea anemones the production of venom occurs also in ectodermal gland cells, which may represent an ancestral secretory pathway for these animals [6].

Within Cnidaria, the Anthozoa is a large, well-supported clade comprising over 6,000 species, and it includes both hexacorals (e.g.: sea anemones and reef-building corals) and octocorals (e.g.: soft corals, sea fans and sea pens) [7,8]. Anthozoa are the subject of active investigation on venoms, but traditionally most studies have been focused on sea anemones (Hexacorallia, Actiniaria) [9–15]. Despite a reduced taxon sampling, these studies highlighted Actiniaria produce complex venoms, including over 15 different protein families with neurotoxic or cytolytic action [14–16]. Comparative analyses have demonstrated that sea anemones produce a neurotoxin-rich venom, while the venom of jellyfish (Scyphozoa) and hydroids (Hydrozoa) is abundant in cytolysins and enzymes [17]. These neurotoxins have been used since the 1970s to investigate nervous system functioning, and some of them possess a remarkable biomedical potential as drug leads. For instance, a synthetic analogue of the ShK toxin from *Stichodactyla helianthus* (ShK 186, or dalazatide) is in phase II trial for the treatment of several autoimmune diseases [18].

Much less is known about the venoms of Hexacorallia other than sea anemones (*e.g.*, reef-building corals and black corals) and of the other major lineages within Anthozoa: tube anemones (Ceriantharia) and Octocorallia [19]. Venom composition was investigated in four Ceriantharia species only and found predominantly cytolytic [20]. Octocorals, instead, have been completely neglected to date by venom studies, probably since they were supposed to possess few, small, and poorly diversified nematocysts [21]. Only their secondary metabolites have been characterized: these molecules, mostly produced by bacteria of the coral microbiomes, play key roles in defence [22]. Octocorallia include some of the most conspicuous organisms of rocky subtidal communities, both in tropical and temperate seas. These ecosystem engineers play important ecological roles and contribute substantially to benthic communities’ diversity: in the Mediterranean Sea, gorgonians constitute up to 40% of the biomass in rocky sublittoral habitats [23].

The white gorgonian *Eunicella singularis* (Esper, 1791) is widespread and abundant in the north-western Mediterranean [24,25]. Despite hosting *Symbiodinium* zooxanthellae, *E. singularis* displays low levels of primary productivity and relies on heterotrophic nutrition to meet its metabolic needs, feeding mainly on zooplankton integrated with particulated organic matter (POM) [26]. Predation on zooplankton, intraspecific competition and defence from predators (mostly snails, crabs and fish) [27] are physiological activities likely supported, in *E. singularis* and allies, by the production of a specialized venom. Indeed, preliminary evidence have demonstrated that haemolytic peptides are present in different species of octocorals [28], and the diversity and abundance of nematocysts could be higher than previously held [21]. However, toxin diversity has not been addressed in Octocorallia so far, hampering our complete understanding of a crucial eco-evolutionary question concerning Cnidaria biology and evolution: to what extent are main toxin families conserved across Anthozoa? To start answering this question, we provided a first overview of the venom produced by an octocoral, applying to *E. singularis* an integrated proteo-transcriptomic approach, which employed whole colony transcriptomic data to match proteomic data obtained from chemical-induced nematocysts discharge or from the whole body of polyps. Our analyses unveiled several previously unreported putative toxin families, providing the first comprehensive description of the venom of an octocoral species. Leveraging these findings, we aimed to assess the degree of conservation of specific toxin folds of Octocorallia in comparison to other Anthozoa, and to elucidate how venom toxin families diversified throughout the evolutionary radiation of octocorals. Employing a comparative approach on available Octocorallia genomes, we reconstructed the evolutionary patterns of the main toxin families, highlighting distinctive evolutionary innovations specific to Octocorallia. Eventually, by complementing our analyses with untemplated, deep-learning assisted structural modelling of the most relevant toxins, we attempted to infer their function and adaptive value.

## Methods

### (a) Specimens collection, venom sample preparation and RNA and proteins extraction

Colonies of *Eunicella singularis* were collected by scuba diving at a depth of 25-30 m around Giannutri Island, in the Tuscany Archipelago (Italy) and kept alive in seawater until further processing. Polyps were dissected from the colony under a stereomicroscope and pooled in three aliquots of about 20 each (about 15 mg of tissue) that were preserved in Trizol at -80 °C until further processing. Total RNA was extracted using Trizol (Thermo Fisher Scientific), following the manufacturer’s instructions, obtaining 5-10,000 ng with RQN values between 6.7 and 8.8. Proteins from 10 additional polyps were extracted through homogenization in milliQ water, followed by centrifugation and analysis of the supernatant to produce the whole body proteome. Additionally, 10 live colony fragments about 25 mm long were immersed for 30 seconds into 1 mL of EtOH in a 1.5 mL tube to induce nematocyst discharge [29]. Samples were freeze-dried and resuspended in 100 μL milliQ water for proteomic analysis.

### (b) RNASeq

To account for variations in venom expression among samples, three cDNA libraries for RNASeq were prepared using an Illumina TruSeq Stranded mRNA Sample Preparation kit according to manufacturer’s instructions and validated qualitatively and quantitatively on a Fragment Analyser, as well as by qPCR on a ROCHE LightCycler 480. The three libraries were then processed for paired-end sequencing-by-synthesis in an Illumina HiSeq 2500 at the MGX facility (Montpellier, France), and quality controlled using FastQC. After trimming, we obtained a number of reads longer than 30 bp and quality above Q20 comprised between 21 and 30 million.

### (c) Bioinformatic analyses of transcriptomic data

Paired-end raw trimmed sequencing data obtained from the three libraries were pooled and *de novo* assembled using Trinity v.2.5.1 [30] with default settings, allowing a minimum contig length of 200 nucleotides. The assembled transcriptome was virtually translated using TransDecoder v.5.01 (https://github.com/TransDecoder/), setting the minimum allowed ORF length to 100 codons.

Expression levels of toxin transcripts were calculated based on unique gene counts as Transcripts Per Million (TPM), as this normalised unit allows an efficient and reliable comparison of gene expression levels both within and between samples [31]. Briefly, trimmed reads from each library were separately mapped against the transcriptome assembly, previously filtered to remove sequence redundancy by considering only the longest transcript for each gene model.

### (d) Protein digestion

Protein extracts were denatured, reduced and alkylated. Briefly, the volume of each sample was adjusted to 100 µL with 100 mM triethylammonium bicarbonate (TEABC). 1M DTT (1 μL) was added to each sample, which was then incubated 30 min at 60 °C before adding 10 μL 0.5 M Iodoacetoamide (IAA). Alkylation lasted 30 min in the dark. Enzymatic digestion was performed by adding 2 μg trypsin (Gold, Promega, Madison USA) in TEABC 100 mM and incubating overnight at 30 °C. Then, peptides were purified and concentrated using OMIX Tips C18 reverse-phase resin (Agilent Technologies Inc.) according to the manufacturer’s specifications. Peptides samples were finally dehydrated in a vacuum centrifuge.

### (e) Shotgun proteomics

Samples were resuspended in 20 μL of 0.1% formic acid (buffer A) and 1 µL was loaded onto an analytical 50 cm reversed-phase column (0.75 mm i.d., Acclaim Pepmap 100® C18, Thermo Fisher Scientific) and separated with an Ultimate 3000 RSLC system (Thermo Fisher Scientific) coupled to a Q Exactive HF (Thermo Fisher Scientific) *via* a nano-electrospray source, using a 120-min gradient of 2 to 40% of buffer B (80% ACN, 0.1% formic acid) and a flow rate of 300 nL/min. MS/MS analyses were performed in a data-dependent mode. Full scans (375 – 1,500 m/z) were acquired in the Orbitrap mass analyser with a 60,000 resolution at 200 m/z. For the full scans, 3 x 10^6^ ions were accumulated within a maximum injection time of 60 ms and detected in the Orbitrap analyser. The twelve most intense ions with charge states ≥ 2 were sequentially isolated to a target value of 1 x 10^5^ with a maximum injection time of 45 ms and fragmented by HCD (Higher-energy collisional dissociation) in the collision cell (normalised collision energy of 28%) and detected in the Orbitrap analyser at a resolution of 30,000.

### (f) Bioinformatic analyses of proteomic data

PEAKS Studio 8.5 (Bioinformatics solutions, Waterloo, ON, Canada), a de novo assisted database software [32-34] was used to analyse MS/MS data from *E. singularis* venom, with parameters set as in [35]. The MS/MS spectra obtained from shotgun proteomics were matched to a custom database resulting from the whole-body transcriptome of the same colony, translated in the six open reading frames.

Protein isoforms were considered only when their existence was supported by MS/MS fragmentation data, whereas translated transcripts identified by PEAKS Studio as belonging to the same protein group were clustered with CD-HIT [36] using an identity threshold of 90%. Only the most representative sequence was retained for the subsequent annotation.

Protein annotation was performed as detailed in the Suppl. Met. and Fig.1. Toxin nomenclature of mature polypeptides followed Oliveira et al. [37]

**Figure 1.**
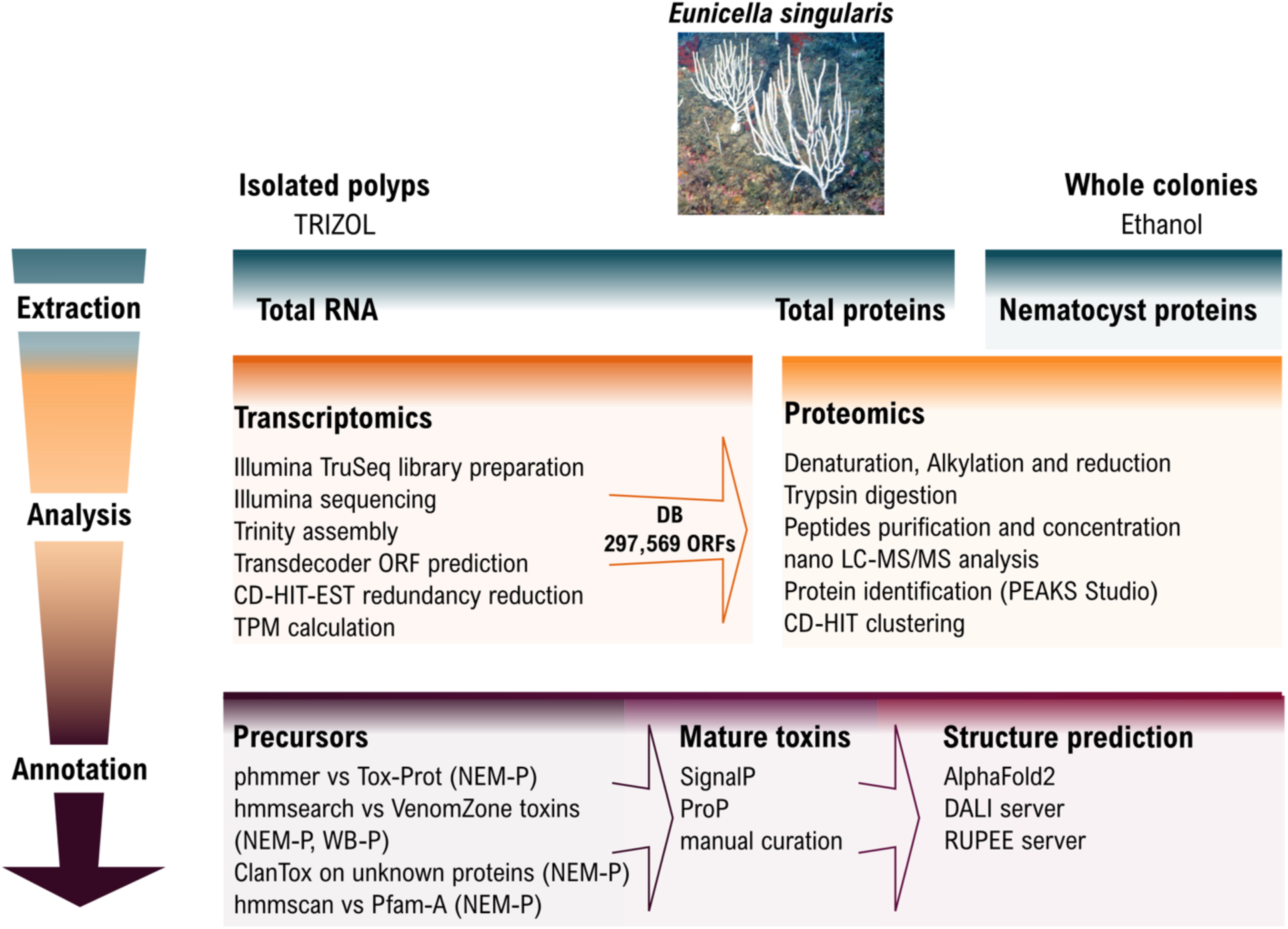
Schematic representation of proteotranscriptomics and bioinformatic annotation pipelines employed in this work.

Structure predictions of mature toxins were performed with the colabfold implementation of AlphaFold2, using MMSeqs2 for multiple sequence alignment generation [38,39]. To increase model confidence for the most challenging targets, custom multiple sequence alignments with orthologue octocorallian sequences were used as inputs and each model was built using 12 recycles. All predictions were performed without the use of templates and energy minimization. The structures that were modelled with reasonable confidence (average pLDDT ≥ 70, which is generally recognised as the threshold for reliable models) [40] were submitted to the DALI [41] and RUPEE [42] servers for structural homology search against the Protein Data Bank. Homology modelling of the saposin-like toxins in the closed conformation was performed with Modeller v9.24 [43]. Electrostatic surfaces were calculated with APBS [44]. UCSF ChimeraX was used for model analysis, visualization and graphics [45].

### (g) Comparative genomic analyses

Toxins identified in *Eunicella singularis* were used as queries for sequence homology searches against a selection of available anthozoan genomes, representative of the diversity of Octocorallia [46] and major Hexacorallia lineages summarized in Suppl. Tab. 1.

Sequence homology searches were carried out with tBLAStn [47], based on an initial E-value significance threshold arbitrarily set to 10. Potential hits were manually curated, by annotating the inferred CDS, detecting putative splicing donor and acceptor sites with the aid of Genie [48]. The presence of *bona fide* orthologues between *E. singularis* and the other species was assessed through combined observations, i.e. (i) the conservation of the main topological feature of the encoded precursor proteins, such as the identification of a signal peptide and the cysteine residues involved in the formation of the disulphide bonds (whenever present); (ii) the shared presence of conserved protein domains, detected with a HMM profile approach, based on the Pfam-A database; (iii) the conservation of gene architecture (e.g. intron number, phase and position), with respect to the orthologous gene from the congeneric species *E. verrucosa*. Whenever needed (i.e., U-GRTX-Esi20 and 21), a fine-scale classification of orthologous sequences was aided by Maximum Likelihood phylogenetic inference analyses, carried out with IQ-Tree [49] with 1000 ultrafast bootstrap replicates based on the best-fitting model of molecular evolution detected by ModelFinder [50].

## RESULTS AND DISCUSSION

### (a) *E. singularis* produces a complex venom

The extraction of venom from cnidarians, like *E. singularis*, is performed by stimulating the animals to fire their nematocysts and then purifying the toxins. Traditional methods use electrical or mechanical stimulation followed by toxin isolation from seawater. Alternatively, nematocysts can be isolated by density gradient centrifugation and mechanically disrupted to release their contents. Electrical stimulation of *E. singularis* colonies proved insufficient to recover enough crude venom for subsequent analysis. Hence, we attempted an ethanol-induced discharge method. While this has been validated for the box jellyfish *Chironex fleckeri* [29], its use for Anthozoan venoms has never been reported. Our results showed that the protein fraction recovered from ethanol-immersed *E. singularis* colonies had a composition comparable to the contents of nematocysts: proteomic analysis confirmed 162 translated transcripts (NEM-P, Suppl. Tab. 2), reduced to 108 non-redundant precursor protein sequences. Based on an extensive bioinformatic annotation (see Methods, Suppl. Met. and Fig. 1), 33 of these sequences were associated to structural components or basal physiology functions, 15 were enzymes, 28 were classified as unknown and 32 were identified as putative toxins (Fig. 2B).

**Figure 2.**
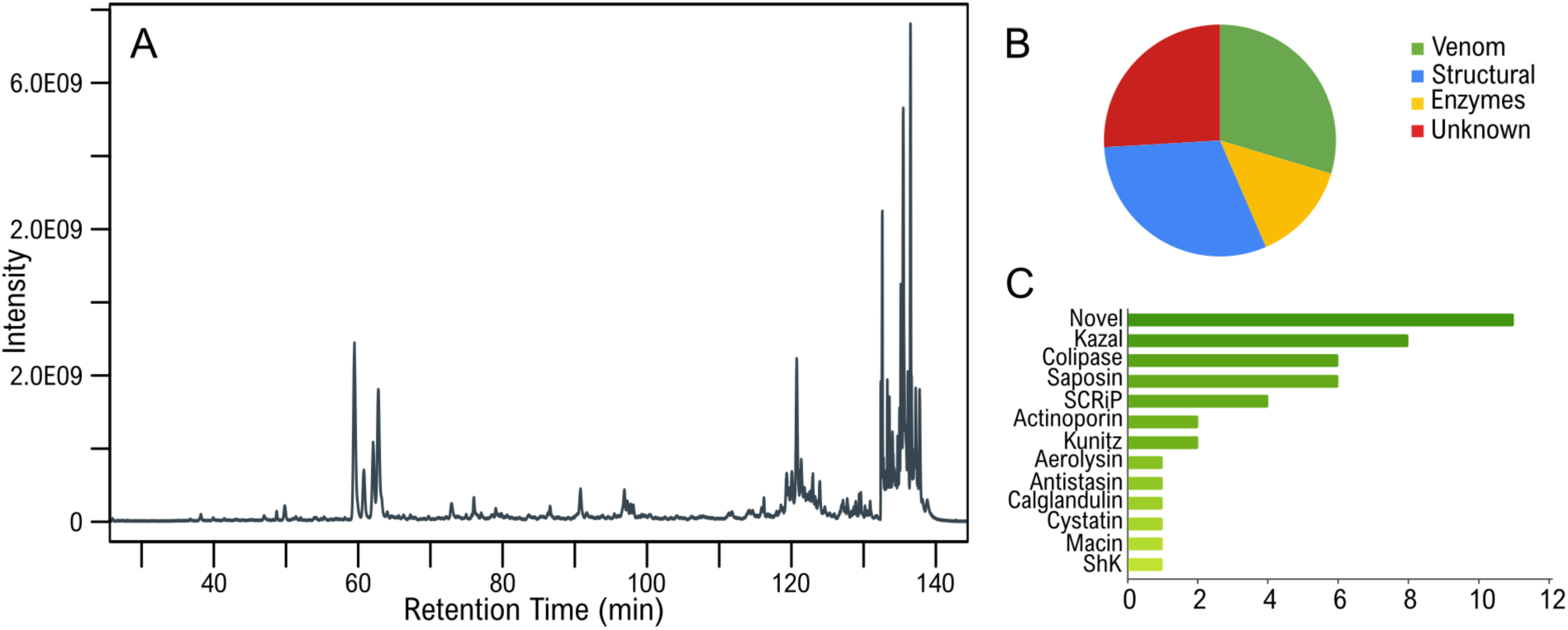
Structural and functional annotation of the venom from *E. singularis.* (A) Total Ion Chromatogram (TIC) of the ethanol extract; (B) Functional annotation of the translated transcripts confirmed at the protein level in the NEM-P; (C) Fold distribution of the mature toxins produced by *E. singularis* based on HMM profile search; the x axis represents the number of proteomics-validated sequences.

Only 17 of them could be also detected also in the whole-body proteome (WB-P, Suppl. Tab. 3) of *E. singularis,* which, by comparison, comprised ca. 1400 translated transcripts. As the NEM-P constitutes a subset of the WB-P, its enrichment in putative venom components served as indirect confirmation of nematocysts firing due to the ethanol treatment. To identify additional putative toxins from other tissues (*e.g.*, ectodermal glands), we also performed a targeted HMM search in the WB-P, looking for known cnidarian toxin families, as defined by VenomZone (https://venomzone.expasy.org). This search produced only four sequences previously undetected in the NEM-P: a protein with homology to the Small Cysteine Rich Proteins (SCRiPs) family and three cytolysins (see below). The absence of these proteins in the NEM-P could be due either to incomplete ethanol-induced nematocyst firing due to differential reactivity of diverse types of nematocysts, or to the existence of additional venom delivery systems, such the ectodermal glands, as previously demonstrated for sea anemones [51], that were not affected by ethanol exposure. In alternative, these proteins may lack a venom function despite having toxin-like folds (e.g. they might be involved in neurotransmission or digestion). The NEM-P comprised some transcripts encoding precursors of multiple mature toxins, which were further classified by predicted fold (Fig. 1C and Suppl. Tab. 4). In total, we identified 44 mature toxins, including a calglandulin, a protein characterized by multiple EF-hand motifs with an accessory role in venom production and processing, omitted from the detailed discussion below.

Following [37], the newly characterized toxins were collectively termed “Gorgotoxins” (GRTX-Esi), according to the most conservative placement of *Eunicella* in the family Gorgoniidae, although recent work has proposed a separate family for this genus (Eunicellidae) [46].

### (b) Established toxin folds are present in the venom of *E. singularis*

#### Small Cystein Rich Proteins-like (octo-SCRiPs)

We retrieved four toxins with homology to hexacorallian SCRiPs, which are known to induce severe neurotoxicity in zebrafish larvae, possess antimicrobial activity and potentiate the transient receptor potential ankyrin 1 (TRPA1) in mouse [52-54]. These comprised U-GRTX-Esi1 (particularly abundant at the transcript level, TPM 1585), 2a-b and 3 (Fig. 3). These proteins exhibited a peptide coverage of the predicted mature sequence exceeding 95%, shared a significant homology and up to 68% pairwise sequence similarity with hexacoral SCRiPs. Despite this similarity, each toxin showcased distinct features.

**Figure 3.**
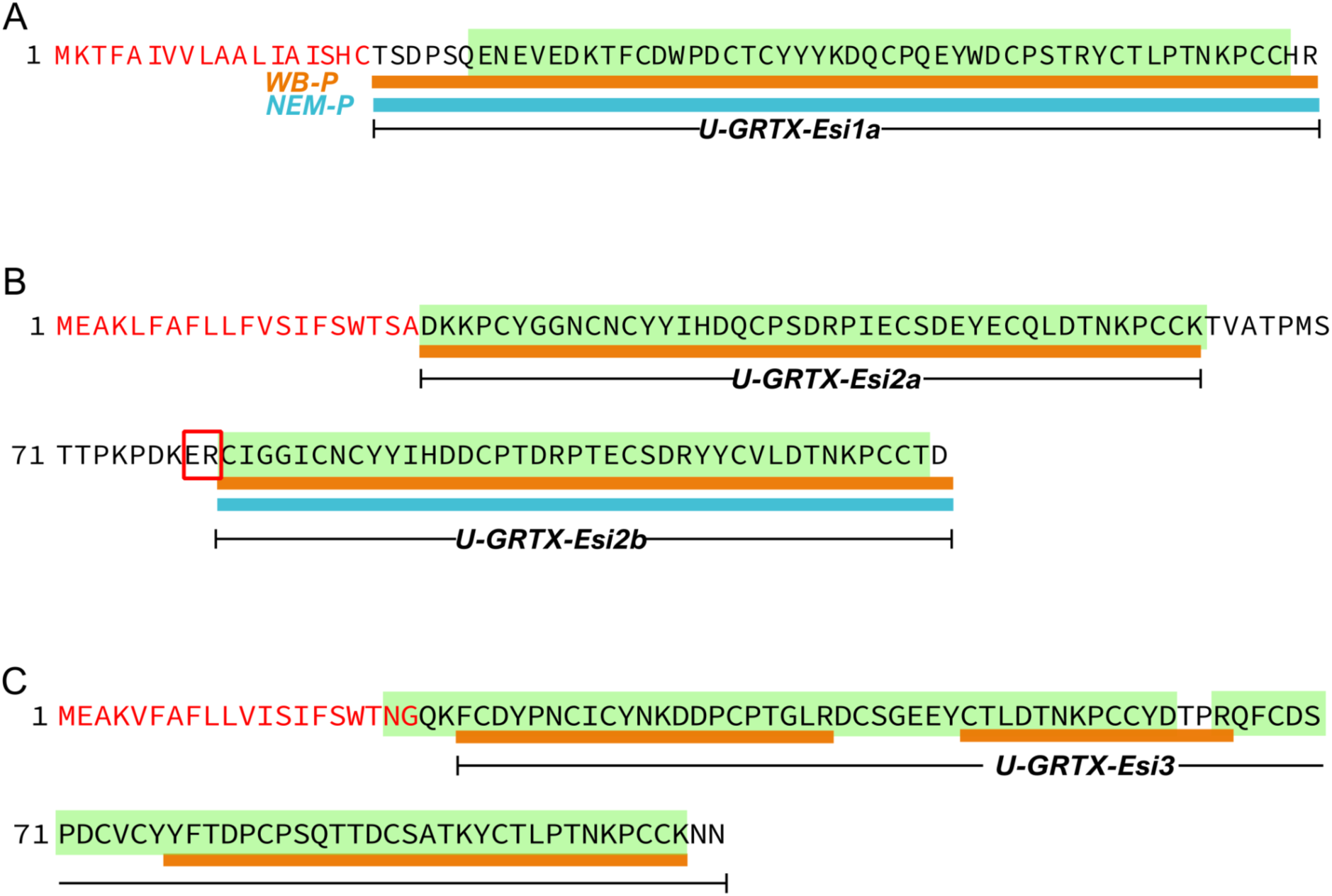
Tissue-specific maturation patterns of the SCRiP-like toxins- from *E. singularis.* The sequences of the precursors of U-GRTX-Esi1 (A), 2a-b (B) and 3 (C) are reported. Red residues correspond to predicted signal peptide, green boxes define the SCRiP domains, the red rectangle defines a PQM protease recognition site [56], orange and blue lines indicate the peptide coverage of the precursors in the WB-P and NEM-P, respectively.

In particular, U-GRTX-Esi2a-b derived from a precursor containing two consecutive highly conserved SCRiP-like domains (pairwise identity 78.9%), separated by a protease cleavage site. Interestingly, while both forms could be detected in the WB-P, only U-GRTX-Esi2b was detectable in the nematocyst discharge (Fig. 3B and Suppl. Fig. 1).

U-GRTX-Esi3, solely detectable in the WB-P, featured a more complex architecture, comprising two SCRiP-like domains in the mature polypeptide, confirmed by the proteomic coverage (Fig. 3C). The absence of predicted cleavage sites between these domains would qualify U-GRTX-Esi3 as a Secreted Cysteine-Rich Repeat Protein (SCREP), an emerging typology of venom proteins found in various organisms, often with high affinity and avidity towards their biological target and a multi-modal action [55].

Although SCRiPs-encoding genes were identified, often in large numbers and associated with tandemly duplicated gene clusters, in the genomes of all the octocoral species investigated in this study, they have a patchy occurrence in Hexacorallia, supporting a complex evolutionary history characterized by lineage-specific expansions and rapid diversification [55]. It is unclear whether SCREPs with two consecutive SCRiP domains were already present in the latest common ancestor of all Anthozoans, or independently acquired in different lineages. Indeed, clear orthologues of U-GRTX-Esi3 could be identified only in *P. clavata, Trachytela* sp. and *R. reniformis*: this architecture had been previously identified also in the hexacoral *Orbicella faveolata* [53]. However, while the only SCRiP so far reported in Octocorallia (in *Scleronephthya gracillima*) presented a conventional cysteine framework [55], the domains here identified display a shift in the cysteine pattern, which nonetheless did not affect the overall protein fold (Suppl. Fig. 2).

#### ShK-containing toxins

The ShK toxins, first identified in the venom of the sea anemone *Stichodactyla helianthus*, are potent KV1 channels blockers [57]. Our search for ShK domains in the proteomes of *E. singularis* led to the identification of U-GRTX-Esi4 in the NEM-P (TPM 103), with three consecutive ShK domains separated by short linker regions. Only the C-terminal domain has the typical ShKT cysteine pattern, whereas the first two domains present an unusual shift of the C-terminal cysteine. None of the domains of U-GRTX-Esi4 presents the key Lys residue necessary for binding KV1.2 and KV1.3, while the subsequent Tyr residue, also important for binding KV1, is extremely conserved (Suppl. Fig. S3). This architecture resembles the precursors of the NEP3 protein family from *N. vectensis* [58]. However, only the first ShK domain of NEP3 is retained in the mature NEP3 toxin, while U-GRTX-Esi4 lacks putative cleavage sites and we found proteomic evidence for the first and the last ShK domain, suggesting the occurrence of a multidomain protein in *E. singularis*.

#### Protease inhibitors

Toxins containing domains associated with protease inhibition are widespread across the tree of life. Among them, Kunitz-like peptides are frequently found in venoms [59] where they can act as inhibitors of several endogenous and exogenous proteases [60] but also show neurotoxic and neuromodulatory activity on a variety of receptors and ion channels [61,62], sometimes with multimodal action [57,59]. Kunitz-like toxins that bind ion channels can be used to paralyze prey and are being extensively studied for their analgesic and neuroprotective properties [63-67]. Our HMM search revealed the occurrence of two Kunitz-like toxins. PI-GRTX-Esi5, detectable in the NEM-P only, presented two Kunitz domains separated only by an Asp-Ile dipeptide, an architecture previously detected in salivary proteins of blood-feeding animals [67,68]. Since orthologous genes sharing the same 3 exons/2 introns organization of PI-GRTX-Esi5 were not detected in Scleralcyonacea, this domain combination most likely represents a recent innovation of the Malacalcyonacea. The second toxin, PI-GRTX-Esi6, with a single Kunitz domain, was detected both in the NEM-P and in the WB-P, and derived from a precursor with a complex architecture, with the Kunitz-like domain followed by three Kazal-type domains, all separated by propeptidase cleavage sites. Also in this case, proteomic evidence indicated a differential maturation pattern of the precursor, with the Kazal-like peptides from this precursor only retrieved in the WB-P (Suppl. Tab. 2). The Kunitz domain of PI-GRTX-Esi5 presented the Arg residue (conventionally “P1”) associated with the trypsin inhibition, whereas its replacement with a Met in both domains of PI-GRTX-Esi6 suggests a chymotrypsin specificity [69]. In addition, PI-GRTX-Esi6 presented a Leu residue aligning with the crucial position for KV1 binding of LmKTT, a scorpion Kunitz-like toxin [70]. These similarities may hint to multimodal activity for PI-GRTX-Esi5, while the lack of key residue conservation in PI-GRTX-Esi6 does not allow inferring any putative function (Suppl. Fig. 4)

Other peptides present in *E. singularis* nematocysts displaying protease inhibitory domains (Kazal-type, cystatines, antistasins, and macins) were detected but did not present novelty elements; they are briefly described in Suppl. Data.

#### Colipases

Finally, a striking feature within the NEM-P of *E. singularis* was the abundance of toxins with a predicted colipase fold (U-GRTX-Esi14-18). Colipases are knottins containing five disulphide bridges, found in spider, snake and tick venoms [71], but to date not in Cnidaria. The prototypical example, MIT-1 [72] interacts through its N-terminal pentapeptide AVITG with prokineticin receptors, stimulating smooth muscle contractions, and exhibiting hyperalgesic effects [73,74]. *E. singularis* colipases lacked the AVITG pentapeptide, a feature shared with the atracotoxins of the spider *Hadronyche versuta,* whose pharmacological target is unknown to date [75].

Apart from the most abundantly expressed protein, U-GRTX-Esi14, all colipases derived from modular precursors containing multiple domains separated by propeptidase cleavage sites. Both U-GRTX-Esi15a-c and U-GRTX-Esi16 were encoded by genes displaying a five exons/four introns architecture (Fig. 4), well-conserved in several orthologues from multiple Octocorallia and Hexacorallia species, pointing out a shared ancient origin with secondary losses in some lineages. U-GRTX-Esi14 and U-GRTX-Esi17 most likely derive from paralogous gene copies, which underwent structural reorganization during evolution. In detail, U-GRTX-Esi17 entirely lost exon 2, which resulted in a shorter precursor protein with an incomplete disulphide pattern. U-GRTX-Esi14 further lost the colipase-fold encoded by exon 3, which was likely reduced and later fused with exon 1. This scenario is supported by the observations that exon 1 in U-GRTX-Esi14 is significantly longer than its paralogues and includes a dibasic propeptidase cleavage site usually present at the 3’end of exon 3. Nevertheless, such remarkable modifications occurred in the latest common ancestor of all Malacalcyonacea, as evidenced by the identification of U-GRTX-Esi14 and U-GRTX-Esi17 orthologues in *D. gigantea*, *M. muricata*, *P. clavata*, *Trachythela* sp. and *P. subtilis* (the latter limited to orthologues of U-GRTX-Esi14).

**Figure 4.**
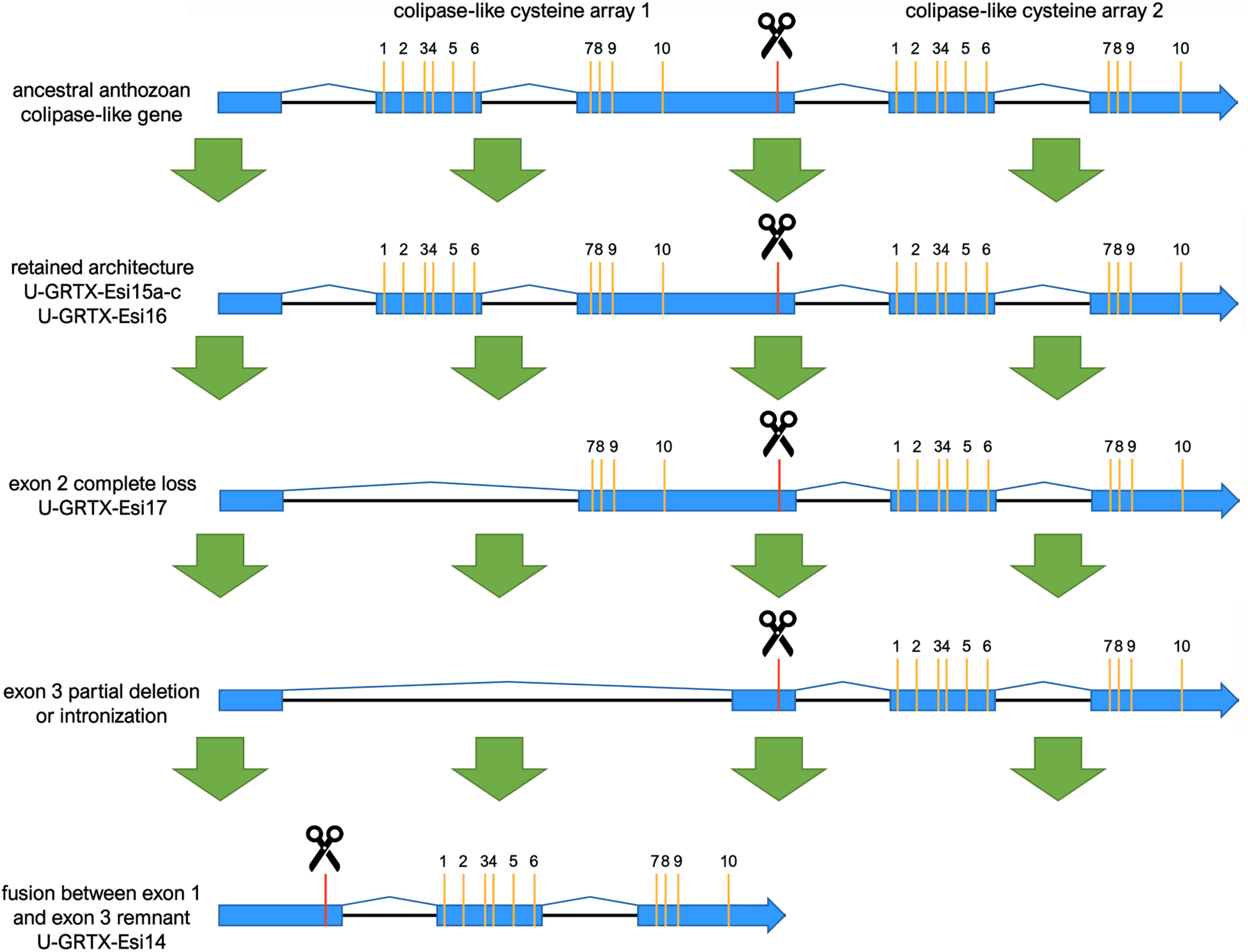
Schematic representation of the evolution of the colipase genes in Anthozoa, with references to specific *E. singularis* genes. Exons are in blue, cysteine residues are indicated by the yellow bars, scissors indicate cleavage sites.

The loss of exon 2 in the gene encoding for U-GRTX-Esi17 results in a shorter precursor protein, with an N-terminal incomplete colipase domain. The maturation of this precursor leads to the release of a polypeptide lacking six out of the ten cysteines (U-GRTX-Esi18), whose presence was also confirmed in the NEM-P. The lack of most of the cysteine scaffold has a major impact on the protein structure: the model predicted with good confidence by AlphaFold2 shows in fact only a long β-hairpin flanked by disulphide stabilized tails.

Notably, the transcripts for all these toxins had exceptionally high TPM values (1806, 569, 826 and 429, respectively for the U-GRTX-Esi14 to 17/18), compared to other proteins detected in the NEM-P (cfr. Suppl. Tab. 4).

### (c) Novel putative toxins were retrieved in the NEM-P of *Eunicella singularis*

We identified in the NEM-P 10 additional proteins possessing the typical sequence features of toxins (small size, abundance of Cys, presence of a signal peptide), lacking sequence similarity with any known protein and often encoded by highly expressed transcripts. Their genomic organization is summarized in Suppl. Fig. 5. High-confidence structural models obtained with Alphafold2 were used to retrieve structural homologues in the Brookheaven Protein Data Bank, in the attempt to infer their biological functions (Fig.5).

**Figure 5.**
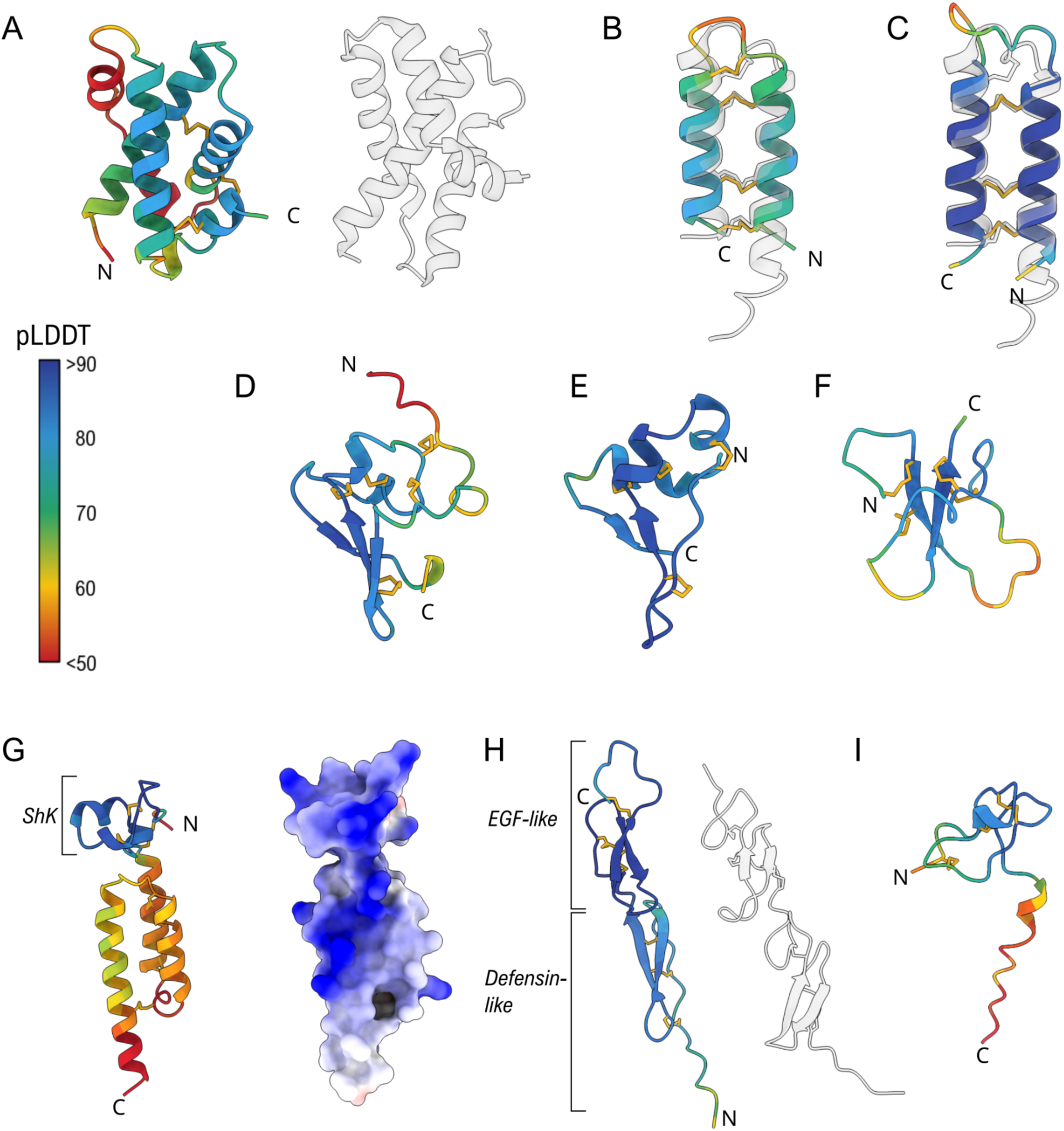
Overview of the structures predicted by AlphaFold2 for the novel toxins from *E. singularis.* Models are coloured according to pLDDT, with higher values (>70) indicating reliable prediction. (A) The putative cytolytic U-GRTX-Esi19 and, on the right, PulS from *K. oxytoca* (PDB 4A56); Overlay of the structures of the two disulphide stabilized helical hairpin toxins U-GRTX-Esi20 (B) and U-GRTX-Esi21 (C) with acrorhagin-1 (PDB 6UX5, right); The predicted structures of the two novel αβ toxins U-GRTX-Esi22 (D) and U-GRTX-Esi23 (E); (F) The SCRiP-like arrangement of U-GRTX-Esi24; (G) The two-domain organization of U-GRTX-Esi25 (left) and its surface charge distribution (right). The ShK domain is adjacent to an all-helical domain. The extended blue patch on the protein surface is common in cytolytic proteins; (H) U-GRTX-Esi26 (left) compared to the byssal protein Pvfp-5β from the mussel *Perna viridis* (PDB 7QAB, right); (I) The structure of U-GRTX-Esi28

Among them, U-GRTX-Esi19 was encoded by an extremely abundant transcript (TPM 2898). The mature protein (100 amino acids, 6 Cys) was detected in both the NEM-P and WB-P and had a weak similarity at the C-terminus with Amoebapore A, the major pore-forming protein of *Entamoeba histolytica*, suggestive of a possible cytolytic function [76]. Most of the protein structure was modeled with high confidence (pLDDT > 80, Fig. 5A), highlighting a compact fold characterized by a knotted four-helix bundle stabilized by three inter-helices disulphide bridges. A similar arrangement, although devoid of disulphide stabilization, is typical of bacterial pilotins PulS and GspS, chaperone lipoproteins that play a role in the assembly of the T2SS secretion system in pathogenic Gram-negative bacteria [77,78]. This putative toxin appears as an evolutionary innovation of Malacalcyonacea, although orthologues are missing in *Phenganax subtilis* and in *Xenia* sp. (Fig. 7, Suppl. Fig. 6). The cysteine framework is always conserved, while the length of some loops varies across species. The abundance of this protein suggests a strong functional relevance, but activity assays will be necessary to clarify its function.

The two putative toxins U-GRTX-Esi20 and 21 did not show any obvious sequence similarity, with a pairwise identity of only 21%. U-GRTX-Esi20 is 48 amino acids long and derives from a longer propeptide proteolytically cleaved at the C-terminus, whereas U-GRTX-Esi21 displays a shorter sequence with a very high cysteine density (8 out of 36 amino acids). The models obtained (Fig. 5B and C) highlighted a structural relationship, as both presented a disulphide-stabilized helical hairpin motif, recently characterized in acrorhagin-1 (U-AITX-Aeq5a) [79], a toxin from the acrorhagi of *A. equina* used in conspecific aggressive encounters [80]. Acrorhagin-1 can induce local tissue necrosis and displays toxicity on crabs, but its mechanism of action has not been clarified, although it has been demonstrated that the toxin does not interact with either potassium or sodium ion channels and it does not possess membranolytic activity [79,80]. U-GRTX-Esi20 and 21 are stabilized by three and four disulphide bridges, respectively, which overlap with those of acrorhagin-1, despite a sequence identity below 20% in both cases.

Our genomic survey indicates that these two toxins belong to two distinct monophyletic orthogroups within a very large superfamily of cysteine-rich peptides, encoded by ancestrally duplicated paralogous genes with intronless structures, that also include other members in *E. singularis* transcriptome, not detected in the NEM-P (Fig. 6). High-confidence orthologues were exclusively found in Malacalcyonacea, except for *P. subtilis* and *Xenia*, forming 6 well-supported clades (Type I to Type VI) (Fig.6). This suggests a rather recent origin of these orthogroups, which derive from a large group of similar peptides also found in other cnidarians, worthy of more extensive investigation. In fact, an ortholog of acrorhagin has been detected in the sensory neurons of *N. vectensis* [81], suggesting an ancestral recruitment from the nervous to the venom system for this class of peptides, followed by lineage-specific diversification in the Octocorallia (Fig. 7).

**Figure 6.**
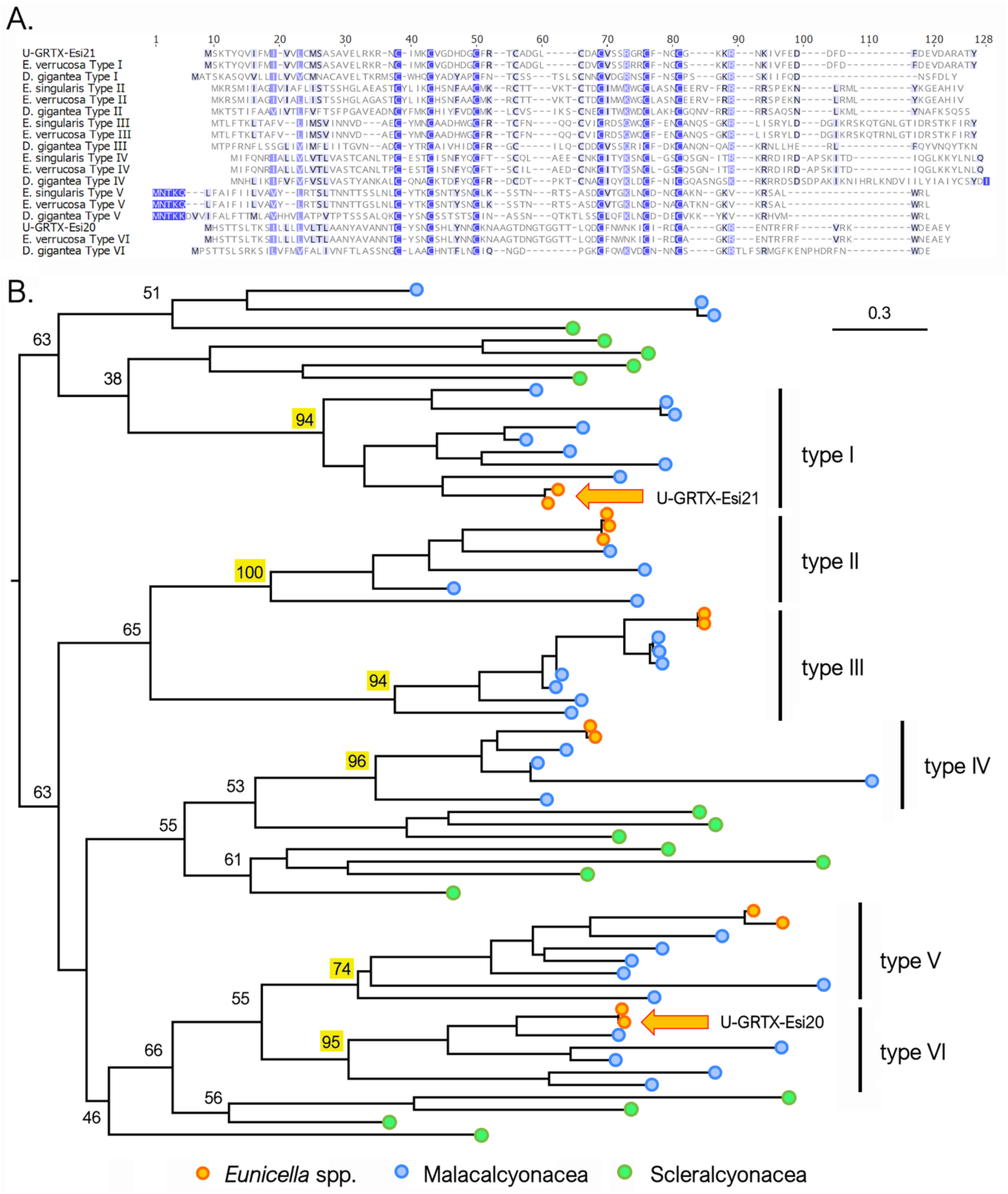
**A)** multiple sequence alignment of the complete precursor amino acid sequences of U-GRTX-Esi20, U-GRTX-Esi21, their paralogs, and their orthologs in *E. verrucosa* and *D. gigantea*. **B)** Maximum likelihood phylogeny of the sequences belonging to the same superfamily of U-GRTX-Esi20 and U-GRTX-Esi21 (indicated with orange arrows), identified in different Malacalcyoneacea and Scleralcyonacea species. All Malacalcyonacea sequences (excluding those from *Xenia*) can be classified within six highly supported monophyletic clades. Bootstrap support values are only shown for the major nodes of the tree, and those denoting the basal node of the six aforementioned clades are highlighted with a yellow background.

**Figure 7.**
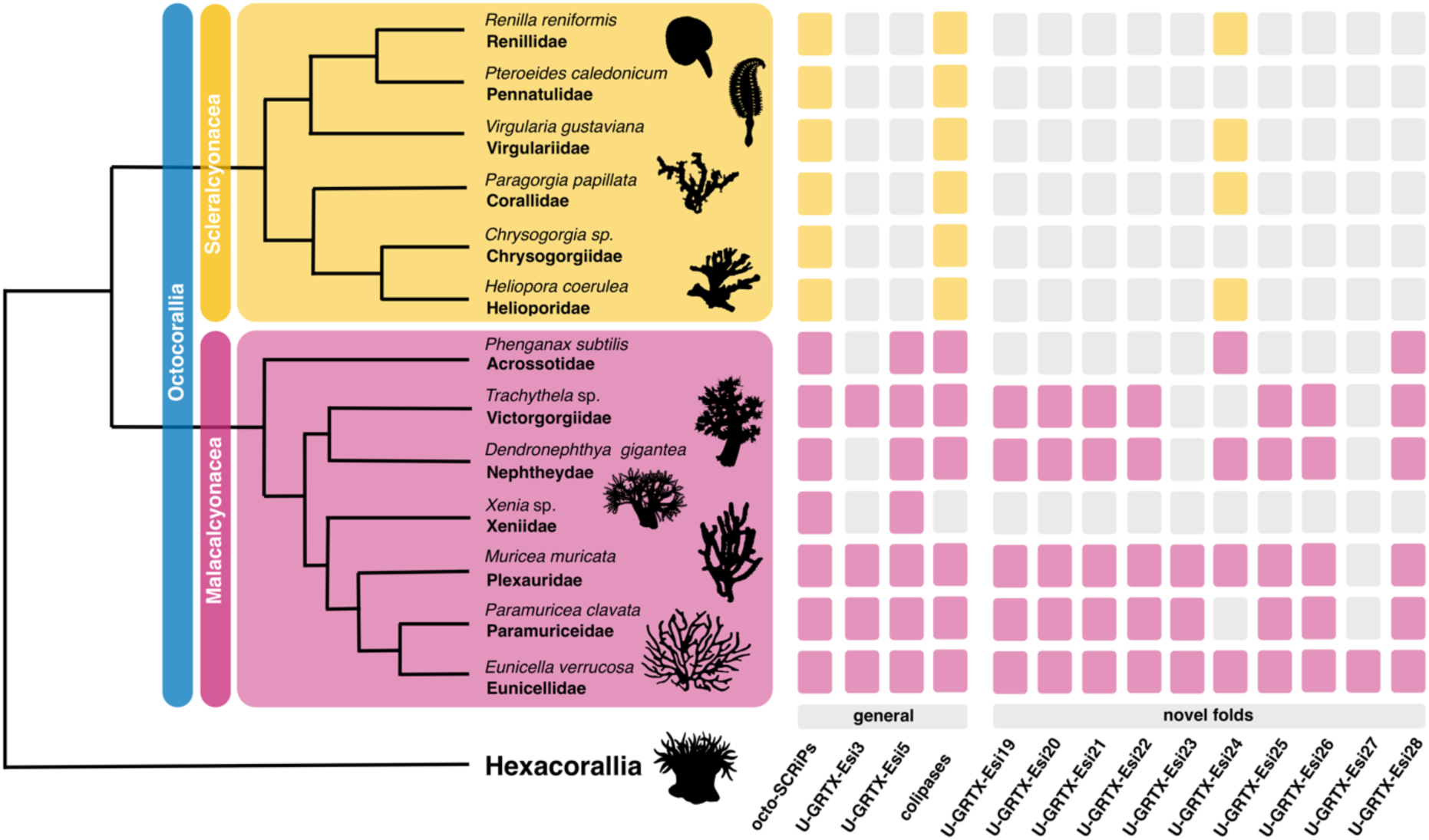
Distribution of orthologues of a subset of *E. singularis* toxins across the Octocorallia genomes here investigated. Phylogenetic relationships of the Octocorallia after [46].

The next two putative toxins, U-GRTX-Esi22 (60 amino acids, 10 cysteines, Fig. 5D) and 23 (45 amino acids, 8 cysteines, Fig. 5E) did not present any sequence similarity with known proteins. In both cases, the models highlighted an unusual disulphide-stabilized αβ fold, suggesting the existence of peculiar and to date undiscovered cnidarian toxin folds. The gene encoding U-GRTX-Esi23 is present with two similar paralogous genes in *E. singularis* and has clearly detectable orthologs in all Malacalcyonacea except *P. subtilis* and *Xenia* sp. (Suppl. Fig. 6, Fig. 7) A divergent paralogue, with a slightly different cysteine pattern, is found in Scleralcyonacea. This toxin most likely constitutes an evolutionary innovation of Malacalcyonacea but could derive from an ancestral sequence with a similar cysteine connectivity present in the latest common ancestor of all Octocorallia. Conversely, U-GRTX-Esi22 presents a single-copy gene, with a detectable orthologue only in *P. clavata* and *M. muricata,* and is probably a recent innovation, only shared by a few related orders within Malacalcyonacea (Suppl. Fig. 7, Fig. 7).

The structure of U-GRTX-Esi24, a 45 amino acids protein with 8 cysteines and a 50% sequence identity with Conotoxin Cltx-2 from *Conus californicus,* could also be modelled with acceptable confidence (Fig. 5F). Despite undetected by HMM profile analysis, the structure presented a SCRiP-like W-shaped fold. This domain has a broad but scattered distribution in Octocorallia, suggesting an evolutionary origin slightly older than most other toxins described in this work. Nevertheless, it is most often found repeated multiple times within the same protein (also in *E. singularis*), with several independent occurrences of single-domain peptides scattered across the tree of life (Fig. 7; Suppl. Fig. 8).

Unique case among the novel toxins from *E. singularis,* U-GRTX-Esi25 was predicted to possess a modular architecture: a recognizable N-terminal ShK domain followed by a region lacking any significant homology. The two domains were clearly distinguishable in the three-dimensional prediction (Fig. 5G), where they appeared as directly connected, without spacer sequences or putative cleavage sites. The unknown C-terminal domain was modelled as an asymmetrical three-helices arrangement, with three disulphide bridges connecting and stabilizing the secondary structure elements. Interestingly, despite the complex architecture predicted, several basic residues from both domains in the model aligned perfectly, forming an extended positively charged patch (Fig. 5G), a feature that suggests a possible membranolytic and/or antimicrobial action [82]. This peculiar combination of domains, paired with a shared 3 exons/2 introns gene architecture, was detected in Malacalcyonacea (except *P. subtilis* and *Xenia* sp). A single orthologous gene was found in *Paragorgia papillata* (Scleralcyonacea), suggesting that the appearance of this protein could predate the split between Malacolcyonacea and Scleralcyonacea (Fig. 7; Suppl. Fig. 9).

The model predicted for U-GRTX-Esi26 (77 residues, 12 Cys) presented an elongated structure made of two domains: an N-terminal region in which two antiparallel beta strands are connected by three disulphide bridges to a proline-rich stretch, in a defensin-like fold found in other cnidarian toxins, and a C-terminal EGF-like domain (Fig. 5H). A similar modular structure was recently described in the byssal adhesive protein Pvfp-5β from the mussel *Perna viridis* [83]; despite the absence of sequence similarity, this structural similarity might indicate for this protein a structural or adhesive function. However, orthologues of the gene were found only in Malacalcyonacea (apart from *Dendronephthya* and *Xenia)*, all sharing a common 3 exons/2 introns gene structure. Three divergent paralogues were found In *P. clavata*, probably originated through a species- or genus-specific gene duplication process (Suppl. Fig. 10).

The last two novel toxins identified in the NEM-P of *E. singularis,* U-GRTX-Esi27 and 28, were short peptides associated with relatively low transcription levels but a high proteomic coverage. In particular, U-GRTX-Esi27 displayed a weak sequence similarity to both SCRiPs and BDS toxins from *A. sulcata*, which are potent ligands of Kv3 channels [84]. The mature sequence of U-GRTX-Esi27, a 39 amino acids long peptide, contained 8 cysteines, 6 of which correspond to the BDS framework. An additional cysteine pair occupied the position of two extremely conserved residues in sea anemone BDS toxins, *i.e.*, a tryptophan and a tyrosine residue, otherwise present in all other members of the family (Suppl. Fig. 11). Only low confidence models could be predicted by AlphaFold2, thus preventing any function inference. Despite the similarity with BDS, orthologous genes encoding for similar proteins were not detected in the genomes of the other Octocorallia considered in this study except from *E. verrucosa.* U-GRTX-Esi27 might therefore constitute an evolutionary innovation of the genus *Eunicella* or the family Eunicellidae (Fig. 7). Finally, U-GRTX-Esi28 possessed a weak sequence homology with the ω-hexatoxin-Hi1g. The modelled structure confirmed this similarity, revealing a cystine knotted core (Fig. 5I), with the addition of an extended and disordered C-terminus, possibly proteolitically cleaved. Orthologues coding for this protein are shared by nearly all Malacalcyonacea but are lacking in *Xenia* (Suppl. Fig. S12). This finding, along with the conservation of the gene structure, in which two exons are spaced by an intron with an equal placement in all species, the signal peptide sequence, and the cleavage sites, strongly indicated that this protein represents an evolutionary innovation of Malacalcyonacea (Fig. 7).

### (d) A diverse range of cytolytic proteins are produced both inside and outside the nematocysts

Our analysis highlighted the absence, in the NEM-P of *E. singularis*, of the common cnidarian cytolytic proteins. A targeted hmmscan search in the WB-P yielded instead positive hits for profiles associated with both the aerolysin (PF01117) and the full anemone cytotoxin (PF06369) families. Coherently with the approach adopted for the NEM-P annotation, we considered only the proteins encoded by complete transcripts presenting a signal peptide; this approach resulted in the identification of three pore forming toxins (PFTs), characterized by diverse structural features. The first one, Δ-GRTX-Esi29, was an aerolysin-like protein, structurally related to the β-PFT Hydralysin [85]; the other two, Δ-GRTX-Esi30 and 31, which shared ∼50% sequence similarity, belonged to the actinoporins family, the most abundant family of cnidarian porins [86]. Interestingly, we observed relevant non-conservative substitutions in *E. singularis* actinoporins, which may indicate selectivity towards different lipids (Suppl. Fig. 13).

Finally, the highest scoring match in the proteomic analysis of the nematocysts content of *E. singularis* corresponded to a 699 amino-acid translated transcript with homology to prosaposin (pSAP) [87]. In humans, the maturation of the orthologous pSAP produces four globular saposins (SAP-A/D), which are co-factors of sphingolipid degrading enzymes. The saposin fold is very ancient and characterizes all the saposin-like proteins (SAPLIPs), which include the amoebapores [88-90]. SAPs and SAPLIPs have a distinctive cysteine framework but display minimal conservation (<25% sequence identity). They are often released in several copies from the same precursor gene [91], assuming a closed, monomeric conformation at lysosomal pH [88,92]. All these proteins are involved in membrane destabilization/solubilization, acting through different mechanisms: lysosomal SAPs undergo a conformational change upon pH shift or in the presence of lipids or detergents, exposing the residues of the hydrophobic core and encapsulating a large number of lipids [93-95]. Other SAPLIPs, like the amoebapores from *E. histolytica* and the caenopores from *C. elegan*s, form oligomeric pores [96,97]. Finally, basic SAPLIPs, like NK-lysin and granulysin, act by “molecular electroporation”, inserting within the membranes [98,99]. In rare cases, the saposin fold has also been detected in non-cytolytic neurotoxins, like the macro-conotoxins con-ikot-ikot and Mu8.1 [100,101], but these proteins are highly dissimilar from SAPLIPs in terms of sequence and cysteine pattern and are encoded by single domain (non pSAP-like) transcripts. The six sequences deriving from the maturation of the pSAP precursor of *E. singularis* (Δ-GRTX-Esi32a-f) were all identified at the proteomic level in the NEM-P. These proteins shared a 24%-65% similarity, higher than what generally observed for homologous SAPLIPs, and were more similar to human saposins SAP-A and C (similarity 26-40%) than to antimicrobial porogenic peptides like amoebapore A (16-25%) or caenopore-5 (13-27%).

We examined the surface potential distribution of the SAPLIPs from *E. singularis*, since the activity of SAPLIPs correlates to their surface electrostatics [102,103]. Except Δ-GRTX-Esi32a and c, they all shared with human SAP-A extended patches of negative charge (Fig. 8), which hints to a pH-dependent mechanism for activation and membrane interaction [104]. In contrast, Δ-GRTX-Esi32a and c presented a more balanced distribution of basic and acidic regions and extended neutral patches, like caenopore-5, suggesting a different, possibly non-monomeric, native arrangement and a pH-independent mechanism of action, possibly through pore formation.

**Figure 8.**
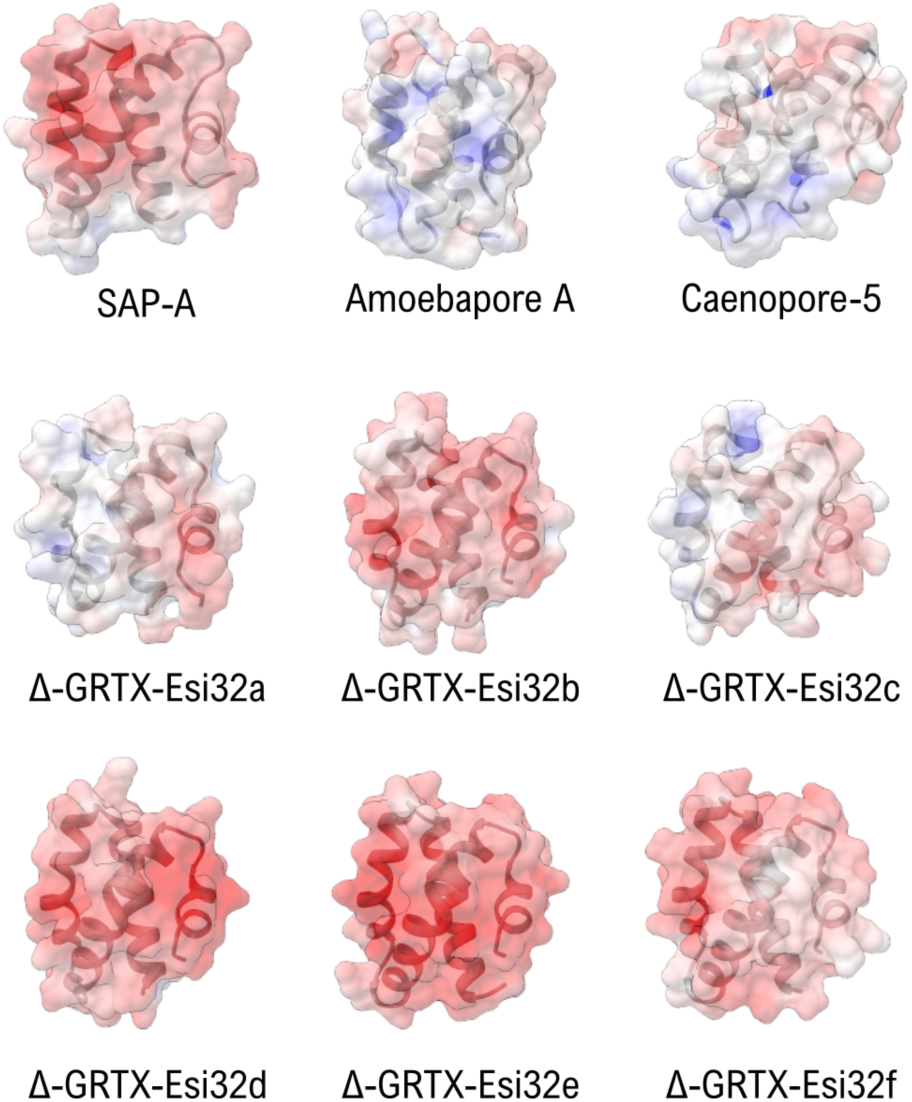
Comparison of the surface electrostatic potentials of the three model saposin-like proteins SAP-A (4UEX), Amoebapore A (1OF9) and Caenopore-5 (2JSA) with the six SAPLIPs identified in the venom of *E. singularis.* The extended patches of negative charge (red) observed in Δ-GRTX-Esi32b, d, e, f are observed in human SAP-A and are indicative of a pH-dependent conformational switch. The lower charge density of Δ-GRTX-Esi32a and c may indicate a different mode of action.

## Conclusions

We have provided the first preliminary depiction of the venomous arsenal of a widespread octocoral. Although experimental limitations did not allow us to conduct our analysis on an isolated venom fraction, by exposing whole polyps of the gorgonian *Eunicella singularis* to ethanol [29] we isolated a sample enriched in proteins presenting typical toxin features. Among them, we detected several folds previously identified in cnidarian venoms, some proteins with structural similarities to venom components of unrelated organisms, and a few with new folds. Even in the absence of functional characterization, the NEM-P seems to provide a good snapshot of the nematocysts content in *E. singularis,* although we could not rule out the possibility that these proteins derive from other tissues. The scarce similarity observed between octocoral and hexacoral putative toxins repertoire is not surprising, given the ancient divergence between these two lineages that recent works dated about 500-700 MYA [2]. In comparison, Elapidae and Viperidae snakes, which present remarkable differences in their venom composition, have diverged only 50 MYA [105]. However, our results clearly indicate, within the NEM-P of *E. singularis,* an elevated content of putative neurotoxic peptides, a feature previously reported for sea anemones. Cytolytic toxins with different structures (and, likely, specificity/function) seem distributed both inside and outside the nematocysts.

The transcriptionally more abundant components of *E. singularis* venom are the potentially cytolytic U-GRTX-Esi19 and the acrorhagin-like U-GRTX-Esi20, the six colipases, the SCRiP-like U-GRTX-Esi1a and the SCREP U-GRTX-Esi4 with three ShK domains. With the exception of the “octo-SCRiPs”, which, however, are characterized by sequence peculiarities, these toxins represent evolutionary innovations of Octocorallia, either shared by the entire lineage (*e.g.,* some colipases), or restricted to the Malacalcyonacea (*e.g.*, U-GRTX-Esi19), or even exclusive of a group of sublineages within the Malacalcyonacea (*e.g.*, U-GRTX-Esi20) (Fig.7).

Putative neurotoxins in *E. singularis* included both novel and established cnidarian folds, the latter often detected within multidomain SCREPs [106]. This is the case, *e.g.,* of the highly expressed U-GRTX-Esi4, with three ShK domains: multidomain ShK proteins have been so far reported in the transcriptomes of parasitic roundworms, in *Hydra* and *Nematostella* [58], and in the vampire snails [107]. The toxin U-GRTX-Esi3 represents another case of SCREP, presenting a double SCRiP domain in the mature polypeptide: this architecture had not been documented in Cnidaria mature toxins to date, although it has been recognized at the transcript level (e.g. in *Orbicella faveolata*). Two domains were also retrieved in the Kunitz BPTI-like PI-GRTX-Esi5, an architecture so far investigated in detail in hematophagous invertebrates only, but extremely abundant among the putative ion-channel impairing SCREP toxins [67,106]. While the abundance of the SCREPs is most likely underestimated across venomous Metazoa, also due to the paucity of proteomic studies confirming the architecture of mature toxins, their abundance in the venom of *Eunicella* might indeed possess some functional significance.

Interestingly, conventional pore forming toxins (PFTs) of Cnidaria, including the aerolysin-like Δ-GRTX-Esi29 and the two actinoporins Δ-GRTX-Esi30 and 31 were not retrieved in the NEM-P. As anticipated, this could depend on the intrinsic limitations of the venom collection procedure employed or reflect an extra-nematocystic production of these proteins. Indeed, finding PFTs outside the nematocysts is not unusual: e.g., hydralysin is an exclusively non-nematocystic PFT involved in prey digestion [84]. Actinoporins, on the other hand, have been to date detected both in the nematocysts and in other tissues. The extra-nematocystic localization retrieved in *E. singularis* might suggest that its actinoporins share the digestive function of the hydralysin-like Δ-GRTX-Esi29. Further histological and functional analysis are needed to confirm this hypothesis, and to evaluate the impact of the observed mutations at critical sites on chemical selectivity and pore formation mechanism of *Eunicella* actinoporins.

Among the proteins of the NEM-P, likely representing the nematocyst content, the cytolytic activity seems to be ascribable mostly to the six saposins Δ-GRTX-Esi32a-f, abundant at the proteomic level. The analysis of their surface charge hints to a considerable diversification of targets and modes of action for this novel group of cytolytic polypeptides: most of them might be inactive at the low pH typical of mature nematocysts but could be activated by the abrupt pH change which follows discharge [108]. On the contrary, the neutral surface electrostatic of Δ-GRTX-Esi32a and c might suggest a pH-independent activation. The novel toxin U- GRTX-Esi19, transcribed at high levels and with a fold compatible with a cytolytic activity, might complement saposins.

While some of these putative toxins are shared by all the investigated octocorals, others had a taxonomic range restricted to Malacalcoynacea and were not retrieved in the analysed Scleralcyonacea, suggesting they might constitute evolutionary innovations arisen after the split between the two major Octocorallia lineages. This is the case of the novel toxins U-GRTX-Esi19 and U-GRTX-Esi28 (Fig. 7). In addition, several toxins were present in a subset of Malacalcyonacea species and are frequently lacking in the representatives of two specific families, Xeniidae (or pumping corals) and Acrossotiidae (i.e., tube corals) (Fig. 7). This peculiar taxonomic distribution might suggest a key role in predation for these toxins, since they are absent in species with a mixotrophic habit, in which autotrophy sustained by symbiotic dinoflagellates is the most relevant source of energy. In contrast, the other Malacalcyonacea included in our analysis (e.g., *Paramuricea*, *Trachytela*) are suspension feeders that largely rely on zooplankton intake [109]. Novel toxins, such as U-GRTX-Esi26, U-GRTX-Esi22, the acrorhagin-like U-GRTX-Esi21 and U-GRTX-Esi27 might thus constitute a predation-related group of putative toxins. In particular, U-GRTX-Esi27 appears to be restricted to *Eunicella* as a genus-specific adaptation (Fig. 7).

The inference on the potential function of octocoral putative toxins is, however, heavily affected by the sparse information available on the biology of these organisms. For example, some Octocorallia species possess sweeper and thread-like tentacles, nematocysts-rich structures that have been hypothesized to mediate interspecific competition, similarly to sea anemones acrorhagi [110]. The occurrence of these structures across the Octocorallia radiation has been poorly investigated to date: this lack of comprehensive information hampers the evaluation of a potential role of specific toxins in competition. Nevertheless, although our comparative genomic approach has revealed a powerful tool to restrict the range of potential functions, further functional characterization studies are needed to unequivocally establish the physiological role of these toxins.

Interestingly, our proteomic analysis pointed towards two cases of differential toxin maturation pathways in the nematocyst vs whole body. The first concerns the two SCRiP isotoxins U-GRTX-Esi2a-b, derived from a double-domain precursor, with both isoforms present in the WB-P while only U-GRTX-Esi2b represented in the NEM-P (Suppl. Fig. 1). The second corresponds to the toxin PI-GRTX-Esi6 and the three isotoxins PI-GRTX-Esi7a-c, encoded by a single transcript in which a Kunitz-like domain (retrieved both in the whole body and in the nematocyst proteomes) is followed by three Kazal-type PI domains only identified in the whole-body proteome. The contribution of alternative cleavage sites to venom diversity has been demonstrated in conopeptides [111], but cell- or tissue-specific differential maturation patterns have not been so far documented for venom toxins, suggesting that the mechanisms regulating venom toxin expression could be more complex than previously assessed.

Through our multidisciplinary approach, we have demonstrated that Octocorallia produce a complex neurotoxin-rich venom, showcasing an overall scarce resemblance to the hexacoral venom repertoire. This distinctiveness is underscored by the presence of novel folds with a complex evolutionary history. These findings call for extensive work, focusing both on the biology of this understudied group of organisms and on the functional characterization of their toxins, to unravel the adaptive significance of Octocorallia arsenal.

## Supporting information

Supplemental References

## Data Availability

The Transcriptome Shotgun Assembly project has been deposited at DDBJ/EMBL/GenBank under the accession GKVI00000000. The version described in this paper is the first version, GKVI01000000. Proteomic data have been submitted to PRIDE with reference 1-20240529-113849-2368759. Other data underlying this article are available in the online supplementary material.

## Acknowledgements

This work has received funding from the European Union’s Horizon 2020 research and innovation programme under the Marie Skłodowska-Curie grant agreement No 748902. The MGX acknowledges financial support from France Génomique National infrastructure, funded as part of “Investissement d’Avenir” program managed by Agence Nationale pour la Recherche (contract ANR-10-INBS-09). Mass spectrometry experiments were carried out using the facilities of the Montpellier Proteomics Platform (PPM, BioCampus Montpellier). Dr. Dany Dominguez-Perez is gratefully acknowledged for his help in data handling and curation.

