## Supplemental Data for "The proteotranscriptomic characterization of venom in the white seafan *Eunicella singularis* elucidates the evolution of Octocorallia arsenal"

**Supplementary Data** - Protease inhibitors

The maturation of the precursor of PI-GRTX-Esi6 produced three Kazal-like peptides, which were designated PI-GRTX-Esi7a-c. Five other proteins containing Kazal-type domains were detected in the NEM-P, namely PIGRTX-Esi8a-b, PI-GRTX-Esi9 and PI-GRTX-Esi10a-b, the latter also characterized by the presence of tandem repeats. Kazal-like protease inhibitors are also present throughout the animal kingdom and abundant components of cnidarian venoms. Notably, Kazal-type peptides have been identified as part of the innate immune system of the cnidarian *Hydra magnipapillata*, where they are produced by the endodermal gland cells and display a potent and selective antimicrobial activity (Augustin et al. 2009). The proteins from *E. singularis* may perform a similar function.

Two more toxins from the NEM-P of *E. singularis* presented domains typical of known protease inhibitors: PIGRTX-Esi11, a cystatine, and PI-GRTX-Esi12, characterized by the presence of multiple antistasin domains. Antistasins are common components of the venom of blood-feeding organisms, such as leeches, where they act as antihemostatic (Iwama et al. 2020). This class of protease inhibitors, detected also in non-blood feeding organisms, had been identified as an endodermal gland-specific product in *Hydra* (Holstein et al. 1992; Hwang et al. 2007).

A protein belonging to the macin family of antimicrobial peptides, U-GRTX-Esi13, could be detected only in the NEM-P of E. singularis. At the structural level, it resembled theromacin from *Theromyzon tessulatum* (Tasiemski et al. 2004), as it displayed an additional Cys pair compared for instance to hydramacin-1 from H. magnipapillata (Jung et al. 2009). In general, macins display marked anti-Gram+ and Gram- activity through a peculiar mechanism of membrane insertion that leads to aggregation and precipitation of bacterial cells (Michalek et al. 2013). In addition, these proteins have been implicated in processes relevant for nerves and tissue regeneration (Schikorski et al. 2008; Jung et al. 2012).
