## Supplemental Figures for "The proteotranscriptomic characterization of venom in the white seafan *Eunicella singularis* elucidates the evolution of Octocorallia arsenal"

### Supplementary Figures

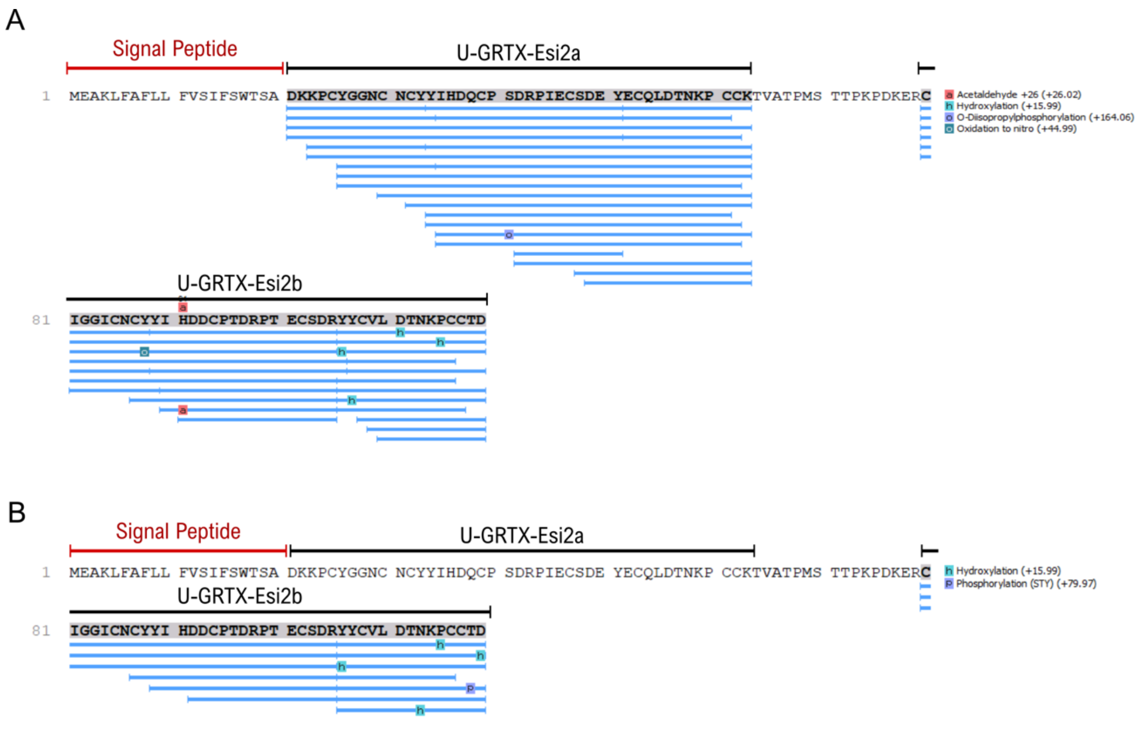


**Supplementary Figure 1.** PEAKS output for U-GRTX-Esi2a/b in **A)** WB-P and **B)** NEM-P. Blue lines below the precursor denote peptides confidently identified by LC MS/MS.

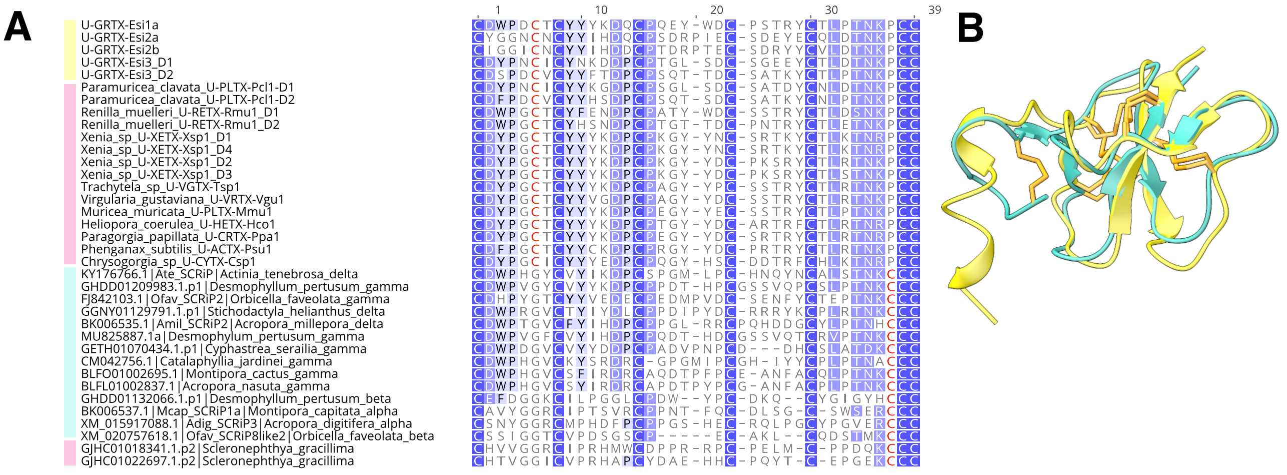


**Supplementary Figure 2. A)** Alignment of the SCRiPS domains of *E. singularis* (yellow bar) with SCRiPS domains retrieved in other Octocorallia (pink bar) and Hexacorallia (cyano bar). Background colour of residues depicts sequence similarity. The cysteine residues whose placement is different in the two main lineages are in red. **B)** Structure overlap of U-GRTX-Esi1 (yellow) and Tau-AnmTx Ueq 12-1 from *Urticina eques* (cyano).

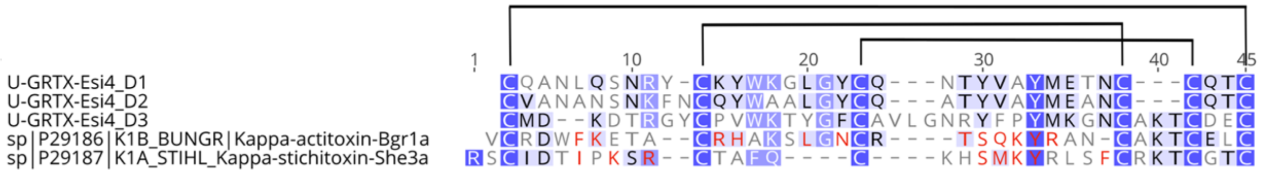


**Supplementary Figure 3.** Alignment of the three domains of *E. singularis* ShK-like protein U-GRTX-Esi4 with the sea anemones mature toxins ShK from *Stychodactyla helianthus* and BgK from *Bunodosoma granuliferum*. Background colour of residues depicts sequence similarity. Residues experimentally demonstrated as relevant for the activity of sea anemone toxins are in red. Disulfide connectivity of sea anemones toxins is also shown.

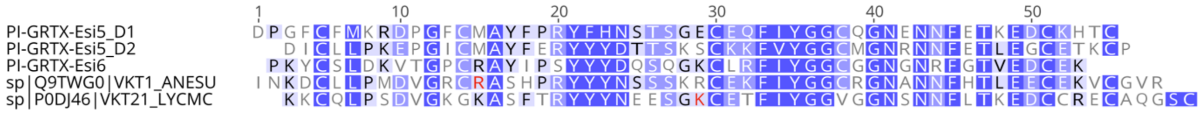


**Supplementary Figure 4**. Alignment of the two Kunitz-type domains of PI-GRTX-Esi5 and PI-GRTX-Esi6 of *E. singularis* with the Kunitz-type serine protease inhibitors KappaPI-actitoxin-Avd3b from *Anemonia viridis* and LmKTT-1a from the scorpion *Lychas mucronatus*. Background colour of residues depicts sequence similarity. Crucial residues for trypsin inhbition are in red.

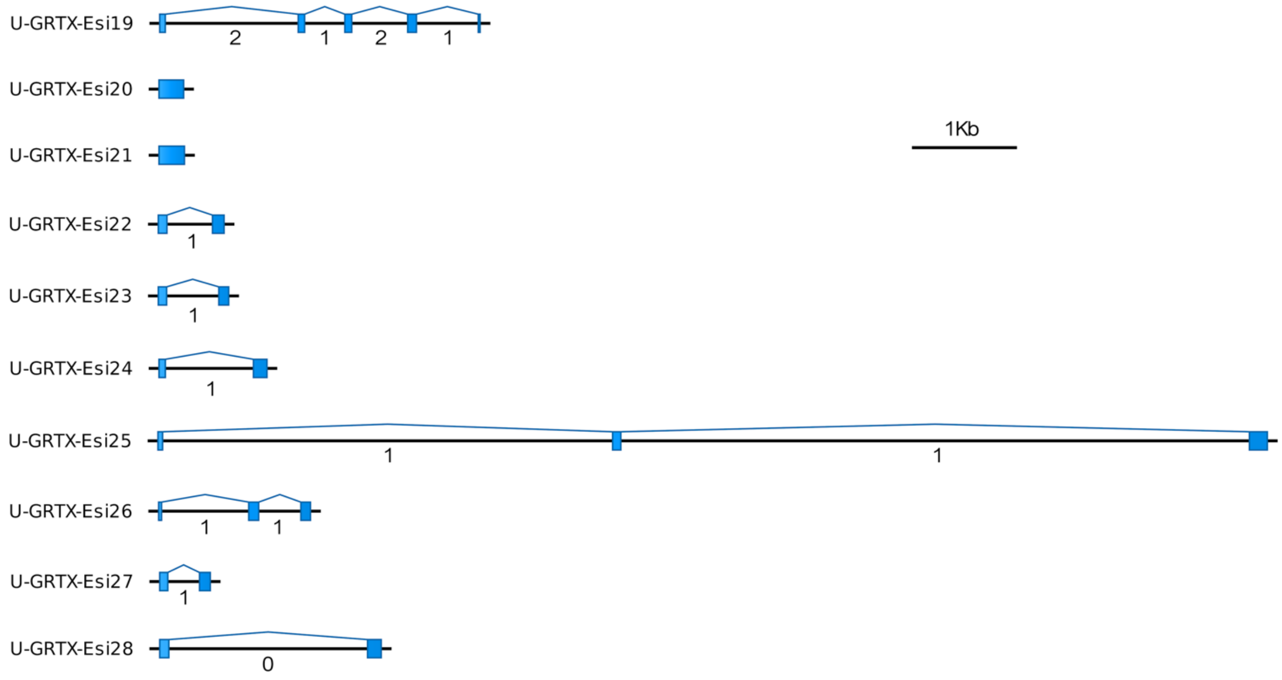


**Supplementary Figure 5.** Genomic organization of the novel toxins identified in *E. singularis*, based on the orthologous genes of *E. verrucosa*. In blue the coding exons.

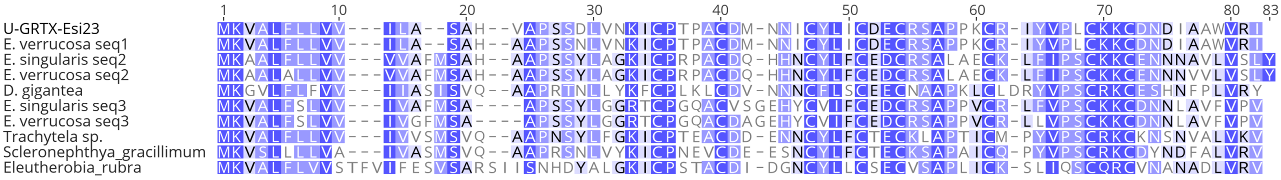


**Supplementary Figure 6.** Alignment of the novel putative toxin U-GRTX-Esi23 from *E. singularis* with orthologs from other Octocorallia. Background colour of residues depicts sequence similarity.

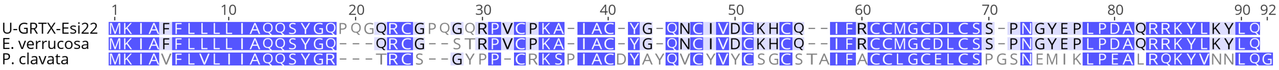


**Supplementary Figure 7.** Alignment of the novel putative toxin U-GRTX-Esi12 from *E. singularis* with orthologs from other Octocorallia. Background colour of residues depicts sequence similarity.

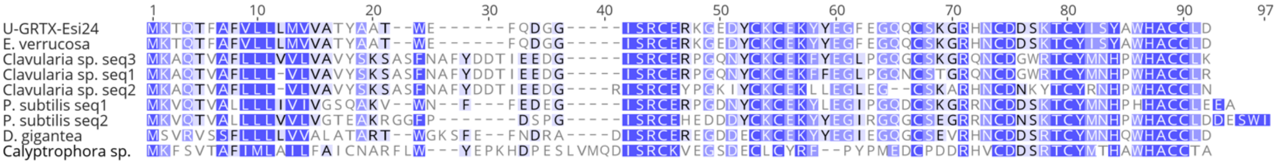


**Supplementary Figure 8.** Alignment of the novel putative toxin U-GRTX-Esi24 from *E. singularis* with orthologs from other Octocorallia. Background colour of residues depicts sequence similarity.

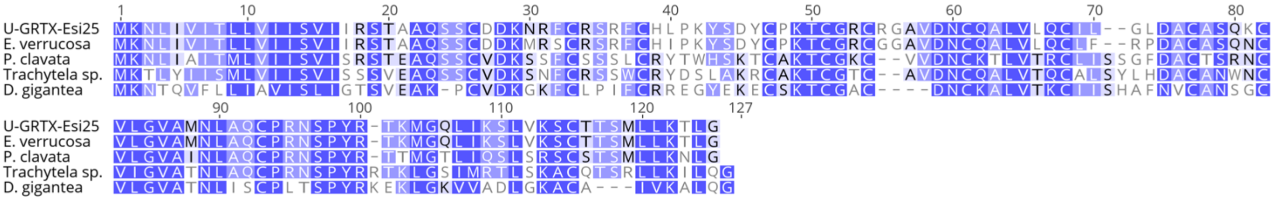


**Supplementary Figure 9.** Alignment of the novel putative toxin U-GRTX-Esi25 from *E. singularis* with orthologs from other Octocorallia. Background colour of residues depicts sequence similarity.

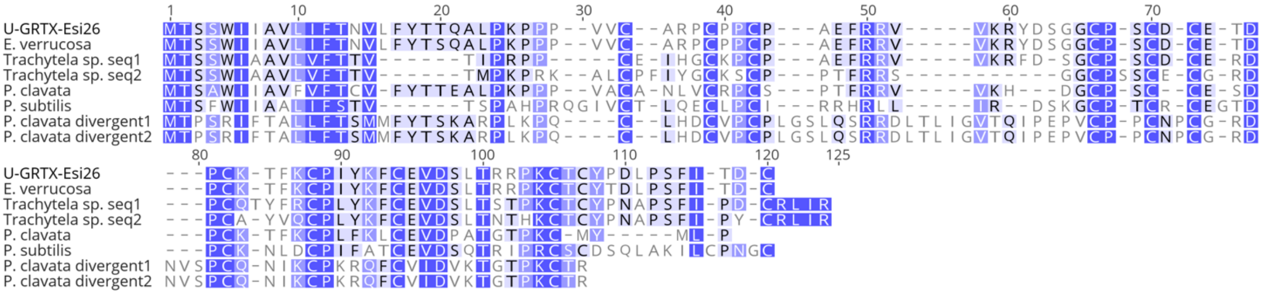


**Supplementary Figure 10.** Alignment of the novel putative toxin U-GRTX-Esi26 from *E. singularis* with orthologs from other Octocorallia. Background colour of residues depicts sequence similarity.

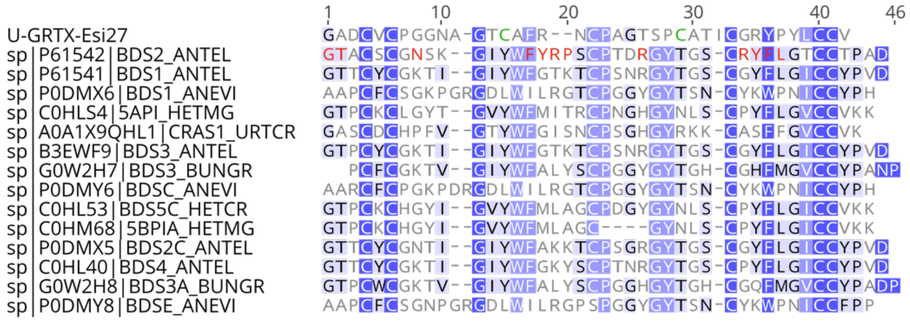


**Supplementary Figure 11.** Alignment of the novel BDS-like putative toxin U-GRTX-Esi27 from *E. singularis* with BDS toxins from several sea anemone species. Background colour of residues depicts sequence similarity. Residues involved in ion channels inhibition are showed in red for BDS2_ANTEL from *Anthopleura elegantissima*. The two additional cysteine residues present in U-GRTX-Esi27 are shown in green.

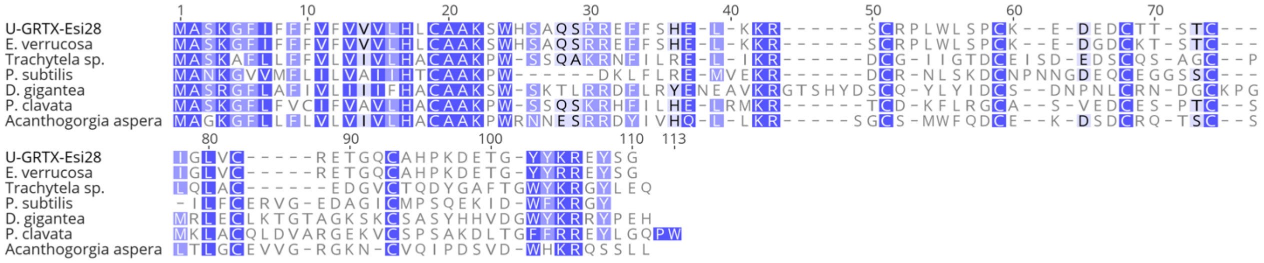


**Supplementary Figure 12.** Alignment of the novel putative toxin U-GRTX-Esi28 from *E. singularis* with orthologs from other Octocorallia. Background colour of residues depicts sequence similarity.

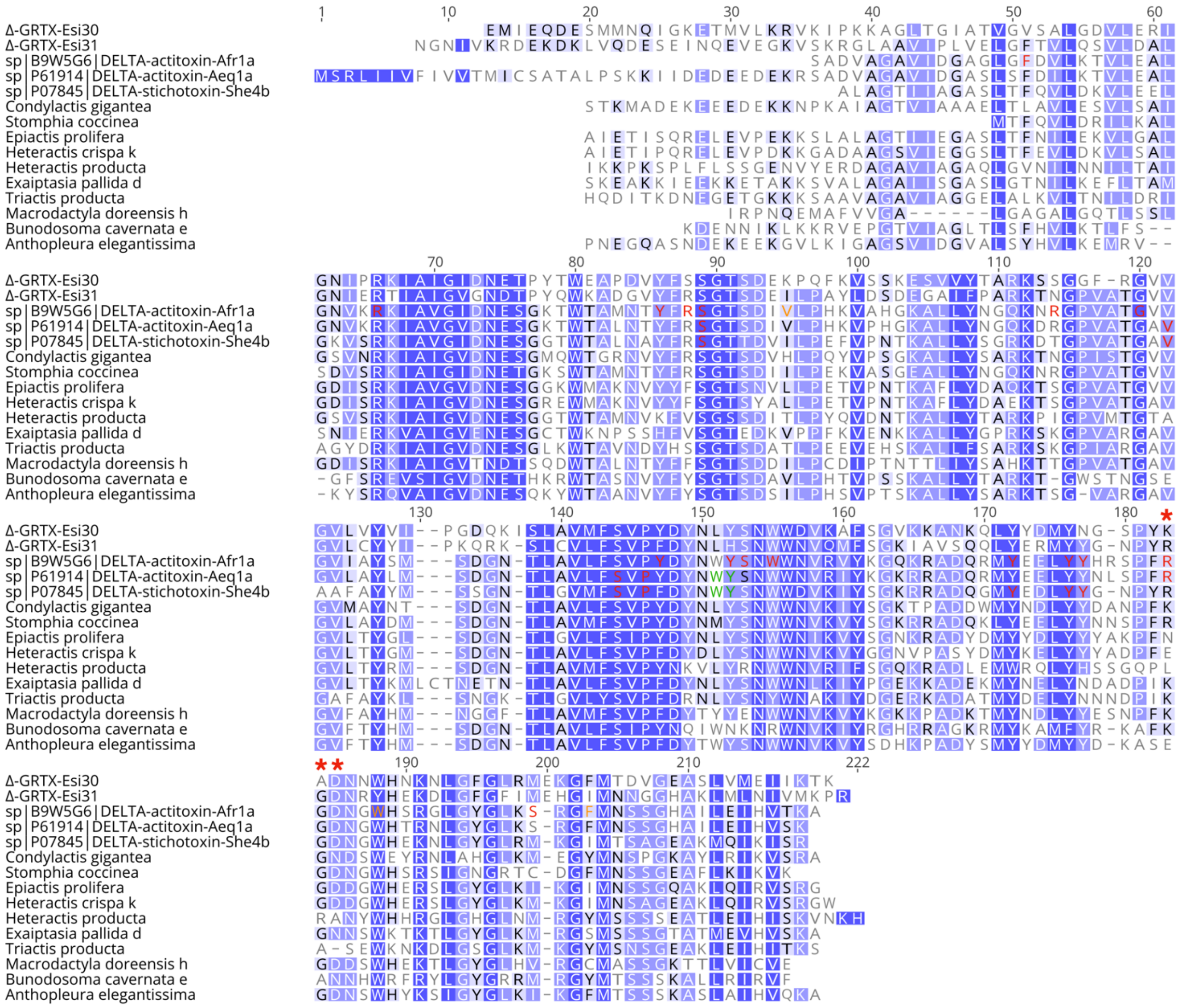


**Supplementary Figure 13.** Alignment of the two *E. singularis* actinoporins U-GRTX-Esi30 and U-GRTX-Esi31 with actinoporins from selected sea anemone sequences from Macrander & Daly, 2016. Background colour of residues depicts sequence similarity. In the functionally characterized toxins from *Actinia fragacea*, *Actinia equina* and *Stychodactyla helianthus*, the residues involved in POC binding are showed in red. The Trp110 and Ty111 determinant in Δ-stichotoxin-She4b for sphingomyelin recognition and initial contact with the membrane, are replaced in Δ-GRTX-Esi30 by a Leu-His dipeptide, suggesting chemical selectivity toward different lipids. In Δ-GRTX-Esi31 the position corresponding to Tyr136 of Δ-stichotoxin- She4b, also in the POC binding site, present an Asn residue, as observed in other sea anemone species. Residues involved in contact with membrane are in green, and residues responsible for oligomerization are in orange. The RGD oligomerization motif, involved in pore formation, is replaced in U-GRTX-Esi30 by a KAD tripepetide (red asterisks); a certain degree of variability in this region, not necessarily affecting activity, is present also in other sea anemone species.
