## Supplemental Methods for "The proteotranscriptomic characterization of venom in the white seafan *Eunicella singularis* elucidates the evolution of Octocorallia arsenal"

**Supplementary Methods** - Toxin annotation pipeline

Protein annotation was performed with HMMER v3.1b2 (www.hmmer.org). A two-step search was conducted on both the whole body and venom proteomes (WB-P and NEM-P, respectively): we first analysed the sequences with the phmmer algorithm against the Tox-prot database (May 2021) and then used hmmsearch to look for specific protein families of the known Cnidarian toxins, as reported by VenomZone (https://venomzone.expasy.org). For this second step, HMM profiles were generated from seed alignments directly downloaded from the Pfam database for the following protein families: Sea anemone cytotoxic protein (PF06369), Aerolysin (PF01117), MAC/Perforin domain (PF01823), *Anemonia sulcata* toxin III family (ATX_III, PF08098), Cystatin (PF00031), Kunitz_BPTI (PF00014), beta defensin (PF00711), ShK (PF01549) and CAP (PF00188). Additional profiles were built with the sequences downloaded from VenomZone for the Cnidaria Small Cysteine-Rich Protein (SCRiP) family, the sea anemone sodium channel inhibitory toxin family, (Type I and Type II subfamilies), the sea anemone structural class 9a family, the sea anemone type 1 potassium channel toxin family (type 1a and 1b), the sea anemone type 3 (BDS) potassium channel toxin family and the sea anemone type 5 potassium channel toxin family. To achieve the most comprehensive annotation possible of *E. singularis* venom, the remaining unknown proteins from the NEM-P were analysed with the ClanTox web server to identify putative novel toxins (Naamati et al. 2009). Finally, a last additional step was performed only on the nematocysts proteome, which was also analysed with hmmscan against the Pfam-A database (May 2021). Signal peptide identification was performed with SignalP v5.0b (Almagro Armenteros et al. 2019) and only sequences corresponding to complete transcripts, comprising a signal peptide and a stop codon, were retained. To identify the sequences of the mature toxins, propeptidase cleavage sites were identified with ProP v1.0 (Duckert et al. 2004) Neuropred (Southey et al. 2006) and by applying the rules described by Kozlov and Grishin (Kozlov and Grishin 2007) and the predictions were validated by comparison with peptide coverage in the PEAKS studio output.
